# *Pinnularia baetica* sp. nov. (Bacillarophyceae): a new diatom species found in an alkaline mountain lagoon in the south of Europe (Granada, Spain)

**DOI:** 10.1101/452110

**Authors:** David Fernádez-Moreno, Ana T. Luís, Pedro M. Sánchezcastillo

**Author notes:** corresponding author’s.

## Abstract

A new benthic freshwater diatom species belonging to the genus *Pinnularia* was found in Laguna Seca of Sierra Seca in the north of the province of Granada, Spain. *Pinnularia baetica* sp.nov. is proposed as a new species based on observations under light (LM) and scanning electron microscopy (SEM) and its special ecology typical of a calcareous lagoon. The most similar taxa to *P. baetica* is *P. atlasii* and with more differences *P. infirma* and the last two were studied through material obtained in lagoons of northern Morocco. Although there are similarities in the morphological characters of the frustule, it was possible to verify through LM and SEM micrographs, evident differences between P. baetica and the other two taxa; on the one end, *P. baetica* has a panduriform shape more pronounced than *P. infirma* and bigger size. On the other hand, the absence of spines in *P. baetica* and the more convergent striation at the poles are the main differences with P. atlasi.

Phylum Ochrophyta Caval.-Sm. (Cavalier-Smith 1995)

Class Bacillariophyceae Haeckel emend. Medlin & Kaczmarska (Medlin & Kaczmarska 2004)

Subclass *Bacillariophycidae* Round (Round et al. 1990)

Order *Naviculales* (Bessey 1907 sensu emend)

Family Pinnulariaceae D.G. Mann, 1990, Genus ***Pinnularia*** C.G. Ehrenberg, 1843

***Pinnularia baetica*** Fernández Moreno & Sánchez Castillo sp. nov

## Introduction

The south of Spain has the richest, most varied and best preserved natural heritage of wetlands in Spain and the European Union. The Andalusian wetlands present a great diversity of ecological types and constitute fundamental areas for the conservation of Biodiversity

The most important studies of benthic lacustrine communities from southern Spain are those of REED (1995), SÁNCHEZ-CASTILLO (1988), SÁNCHEZ CASTILLO (1993), LINARES-CUESTA & SÁNCHEZ-CASTILLO (2007) and SÁNCHEZ-CASTILLO et al. (2008). In addition to LINARES-CUESTA (2003) who made an inventory of the epiphytic and epilithic diatom communities of the lagoons from Sierra Nevada, as well as some works done in Doñana lakes, where some new species were described (BLANCO et al. 2013). Many of the reamaining diatom records were assigned to lothic systems (MARTÍN FARFÁN, 2016).

This older literature does still not reflect all the diatom diversity in the south of Spain, as long as there is not any record in many of the more than 200 wetlands found it in Andalucia.

The species from *Pinnularia* EHRENBERG genus are part of a complex taxonomical group of great diversity, with 708 taxa accepted according to the last revision made in AlgaeBase (GUIRY & GUIRY 2018). Despite of the high diversity of this genus, in the south of Spain is very scarce and the most important populations are in Sierra Nevada Mountain and in acidic rivers as Rio Tinto. Traditionally the species from *Pinnularia* genus have been reported as living in extreme environmental conditions, like the freezing temperatures of the Antartic region where *Pinnularia catenaborealis* (PINSEEL et al. 2016), *Pinnularia sofia* (VAN DE VIJVER et al. 2004), *Pinnularia obaesa* and *Pinnularia australorabenhorstii* (VAN DE VIJVER 2008), and several other species of *Pinnularia* (VAN DE VIJVER & ZIDAROVA 2011; ZIDAROVA et al. 2012) are living well adapted. They can also be present in the surrounding streams of mining areas, characterized by high metal concentrations and very low pH, as it is the case of *Pinnularia aljustrelica* (LUÍS et al. 2012) and *Pinnularia paralange-bertalotii* (FUKUSHIMA et al. 2001). What all of them have in common is the oligotrophic characteristics of their waters (VAN DE VIJVER et al. 2012). Other relevant aspect, is that some of the described species of the Antarctic region can form colonies through the presence of marginal spines (VAN DE VIJVER et al. 2004; PINSEEL et al. 2016), although most of the *Pinnularia* species have isolated life forms (ROUND et al. 1990). Although chain formation by spines affects many biological processes, such as sexual interactions, predation and nutrient uptake (e.g. FRYXELL & MILLER 1978; PAHLOW et al. 1997), it is not clear the main function in other taxa of the *Pinnularia species* (PINSEEL et al. 2016). Among those that have spines it is important to highlight *Pinnularia atlasi* DARLEY (DARLEY 1990). In that study, DARLEY did not show the presence of spines, however in a posterior study carried out in Sardinia by LANGE-BERTALOT et al. (2003), spines were detected in the valvar margin from other population of *P. atlasi*, mostly concentrated in the central area and disappearing towards the ends of the valve.

*Pinnularia infirma* KRAMMER & LANGE-BERTALOT (1985), a second taxon used for comparison was firstly described in central Europe, in a locality situated at 250 m of altitude in Franken, Germany. It was also found in Ouiouane, Morocco, together with *P. atlasi*. No ultrastructural data was provided and consequently, no reference to the presence of spines was given.

This study is part of a larger one that intends to know the diatomological diversity of the coastal zone from the lagoons of Andalusia (southern Spain) (unpublished results).

According to European algal framework directive (POKAINE et al. 2016), studies using diatoms are normally focused in riverine communities and not so much in lacustrine ones. Thus, this is why this study is crucial to improve the knowledge on diatom communities from lagoons, specially alkaline ones.

The main objective of this work is the description of a new species from the genus *Pinnularia*, *Pinnularia baetica* sp. nov. A comparison will be made regarding the frustule morphology of the population found in the province of Granada (South of Spain) both in Light Microscopy (LM) and in Scanning Electron Microscopy (SEM), with *P. atlasi* described by DARLEY, obtained directly from (locotype)the 2 sites of northern Morocco (Ouiouane and Guedrouz) where DARLEY originally obtained the type population. In spite of type material was asked, it was not possible to received. It will also be compared with the description done by LANGE-BERTALOT et al. (2003), in the Island of Sardinia because DARLEY’S description did not include ultrastructural analysis.

## Methods

### Sampling

The epipelic diatom communities of sediments samples came from the littoral zone of three lagoons, one from South Europe (Spain) and two from North Africa (Morocco) (Table 1, Fig. 1):

1. The Laguna Seca (Granada, Spain), that is located at the north of the Natural Park of Sierra de Castril (Complex of Baetic Mountains). It is dominated by outcrops of limestone and dolomites originated in the Upper Jurassic. The Laguna Seca is at 2000 m of altitude, and is small pond located in a shallow basin, filled with rainwater and snowmelt; it is usually dry at the beginning of the dry season. This material was collected in 2007.

**Table 1.** Sampling sites location and its respective collected taxa.

| Site | Locality | Taxa | Coordenates |
| --- | --- | --- | --- |
| Laguna Seca | Granada | *P. baetica* | 37°56'41.86'' N,<br>2°40'54.80'' W |
| Ouiouane | Azrou | *P. atlasi, P. infirma* | 33°07'52.58'' N,<br>5°20'36.72'' W |
| Guedrouz | Marrakech | *P. atlasi* | 31°26'12.45'' N,<br>7°28'06.82'' W |

**Fig. 1.**
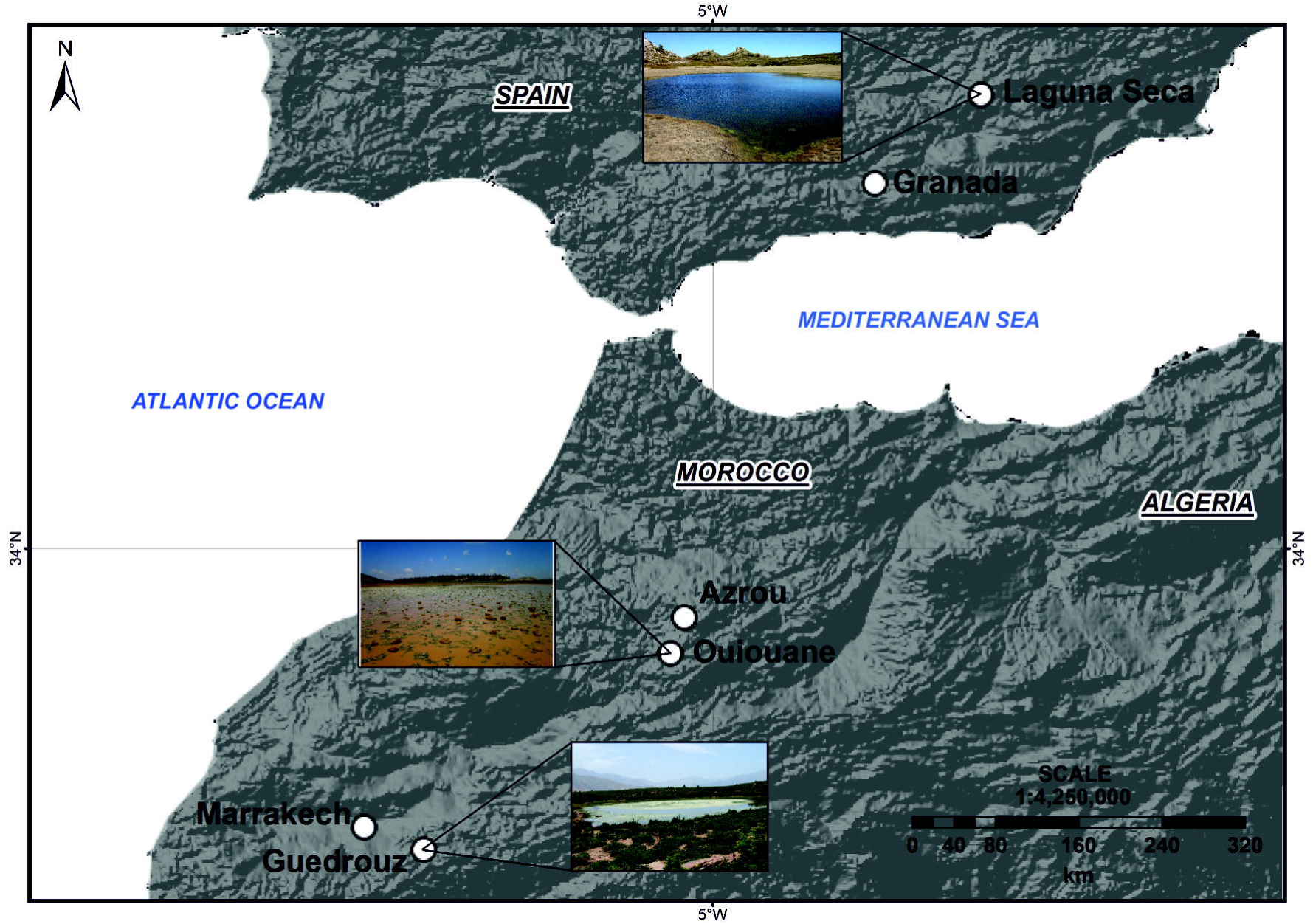
Geographical location of the 3 sampling sites where *Pinnularia baetica* (Laguna Seca), *Pinnularia atlasi* (Ouiouane and Guedrouz) and *Pinnularia infirma* (Ouiouane) were sampled.

The Ouiouane (33°07′52.58′′ N, 5°20′36.72′′ W) and Guedrouz (31°26′12.45′′ N, 7°28′06.82′′ W) were problematic to sampling, because the coordinates were not shown in the original *P. atlasi* described by DARLEY. We did a search on Google earth to find Guedrouz lagoon from the Darley’s description “Timenkar plates, at 2200 m of altitude in the central High Atlas in the region of Marrakech”, finally with the help of people from the high Atlas we found the lagoon. The Ouiouane lagoon was chosen on google earth with a selection of several lagoons close to the city of Azrou at the mountains, as long as Darley described the location close to this city. With the help of field microscope. we distinguished the typical panduriform outline and shape of *Pinnularia atlasi* at the littoral zone of Ouiouane lake, The other lagoons and lakes did not have the presenceof this diatom species. They are temporary ponds that evaporate over a period of 6 to 8 months (DARLEY 1990). Ouiouane is close to the city of Azrou, with an altitude of 1880 m. Guedrouz is located close to the city of Marrakech at about 2000 m of altitude. The material of Guedrouz and Ouiouane were collected in 2014. We proposed this material as a Locotype and it will be submitted it to a collection housed in the Herbarium, University of Granada (Spain).

### Diatoms treatment

Sediment samples were obtained by suction through a one-metre-long glass tube (a Lund tube). Several transepts were made in the littoral zone in order to get an integrated sample of the epipelic community. Field parameters such as: pH, conductivity and temperature were measured *“ in situ”* with a WTW pH/Cond 340i/SET, portable multimeter.

The samples were transported to the laboratory in cold (4°C-8°C) and dark conditions. Once in the laboratory, the pellet was left to stand in a petri dish, then the supernatant was removed and after the sediment was covered by cover slips. After at least 12 hours we proceeded to remove the coverslips where by phototactic movement, the diatoms are attached to the coverslip algae, which were then fixed with 4% formaldehyde.

The samples were oxidized with hydrogen peroxide 30% w/w. Possible calcium carbonate inclusions were removed by adding a few drops of dilute hydrochloric acid 1N into the sample. They were then washed with distilled water four times, leaving the sediment sample and the supernatant was then removed several times to prevent rupture of the valves when the samples were centrifuged. Permanent preparations were mounted with a synthetic resin, Naphrax®, (Brunel Microscopes Ltd, UK) with a high refractive index (1.74). Light microscopy observations were observed using a Leica DMI3000 B light microscope (Phase contrast) with a 100x oil immersion objective with NA (numerical aperture) of 1.30 and LM photographs were taken with a Leica DFC295. Measurements of valve length, width and number of striae (in 10 μm) were taken of at least 20 specimens per species, under the light microscope several subsamples chosen for scanning electron microscopy were filtered through polycarbonate membrane filters with a 5μm pore diameter, mounted on stubs with double sided carbon. A Zeiss SUPRA40VP Electron Microscope with variable pressure high resolution (FESEM) was used for observation of the valve structures. Morphological terminology follows ROUND et al. (1990), KRAMMER (2000) and COX (2004). Apart from the original descriptions of *Pinnularia atlasi* made by DARLEY, the following publications were consulted for taxonomical and ecological comparison: HUSTEDT (1930), KRAMMER & LANGE-BERTALOT (1986, 1988, 1991 a, b), LANGE-BERTALOT et al. (2003), LEVKOV et al. (2005).

After trying to contact with the herbarium of Marrakech, we did not received any reply regarding the type population of *P. atlasi*.

The range of morphometric and micrographic data from *P. atlasi* in DARLEY’s description included both populations of Guedrouz and Ouiouane. In this study also due to the low number of valves of *P. atlasi*, the morphometric and micrographic data were also from both populations of Morocco. For each taxon, the number of valves measured was n=30.

All images were digitally manipulated, and plates were made using CorelDRAW 2017 and the map (Fig. 1) was done using ArcGis vs.10.

## Results & Discussion

### *Pinnularia baetica* FERNÁNDEZ MORENO et SÁNCHEZ CASTILLO sp.nov. Fig. 2-16: LM, Fig. 47-52: SEM

Holotype: Phycotheque GDA-algae slide number 3725, prepared with material from the sample collected in Laguna Seca, housed in the Herbarium, University of Granada (Spain).

Isotype: Slide BM 101 919, prepared with material from the sample collected in Laguna Seca, housed in the Natural History Museum, London (UK).

Type locality: Laguna Seca, Sierra Seca, Parque Natural de la Sierra de Castril, Granada. Spain.

Geographical coordinates: 37º56′41.86′′ N, 2º40′54.80′′ W.

Etymology: The specific epithet *baetica* refers to the name of the mountain chain complex.

### Light microscopy (Figs. 2-16)

Valves panduriform, narrow in the center, then showing a wider enlargement and finishing narrow again close to the poles with cuneate apices. Length 47.2-72.2 µm, width 9.3-12.8 µm. Central area from rhombic (e.g. Fig 6) to broad fascia (e.g. Fig 12), formed by the shortening of one or three central striae. The fascia was observed sometimes on both sides (e.g. Figs 4, 12) or just might be smaller (Fig 5, 6). Axial area linear-lanceolate, widening toward the central area.

The raphe was central, positioned in the middle of the axial area with the proximal raphe endings unilaterally deflected, terminated in slightly expanded but not drop-like pores (Figs 2-16). The distal raphe fissures were hooked. The transapical striae were slightly radiated to parallel and convergent at the poles (9-10/10 µm) (Figs 3-7,11,13,15).

**Fig. 2-16.**
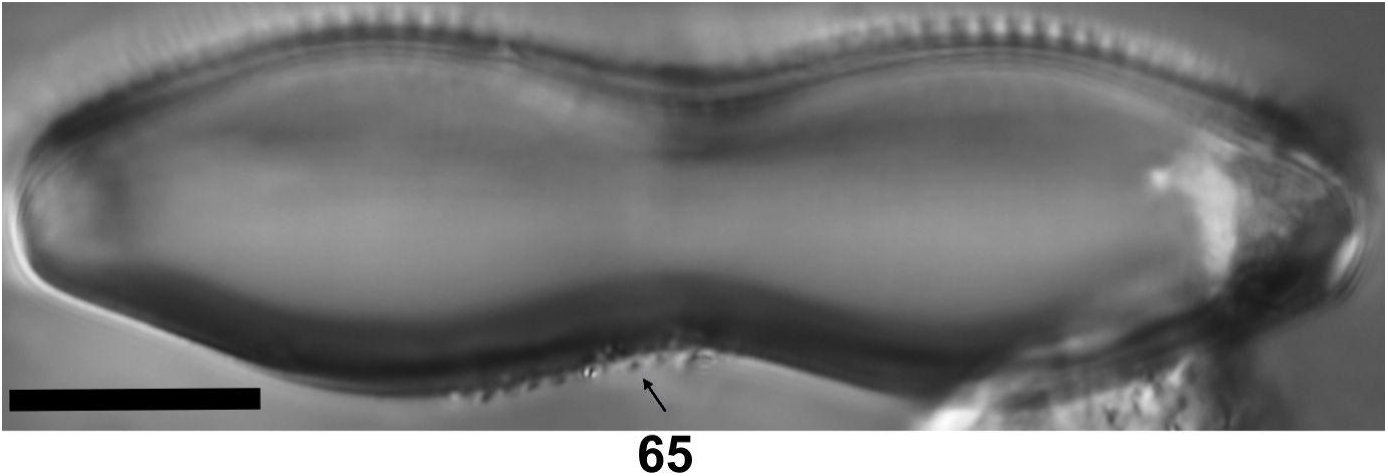
*Pinnularia baetica* sp.nov. Light micrographs of the type population sampled in Laguna Seca in 2007. Scale bar 10 μm.

### Scanning electron microscopy (Figs 48-53)

The external proximal raphe endings were slightly expanded and unilaterally deflected (Figs 47, 51). The distal raphe fissures were clearly hooked shaped (Figs 47, 49), continuing onto the mantle as a curve line downward. The internal raphe was straight with short, bent, proximal raphe endings with a small inflated central nodule (Fig 48). The distal raphe endings terminated onto a small, slightly eccentric, weakly raised helictoglossa (Figs 48, 50). The striae were composed of large alveolus (Fig 50). Externally, each alveolus was composed of 7-8 rows of small areolae (Fig 52) with a maximum diameter of 75-100 nm. Each pore is occluded by a hymen which can be opened by 6/5 sub-pores (20-30 nm diameter) to release the pore surface.

### Ecology

The limnological characteristics of the Laguna Seca at the sampling collection moment (2007), corresponded to the end of the flood period, when the bottom of the pond became full of a community of hydrophilic meadows. The water temperature was 15.7°C, the mineralization was low (83 µS/cm), with a high pH of 9.2. The associated diatom flora from the littoral zone was dominated by *Nitzschia tenuirostris* Manguin (31 %), *Pinnularia* cf. *obscura* Krasske (32 %) and *Pinnularia baetica* Fernández Moreno & Sánchez Castillo sp. nov (12 %) as well as some rare species from *Eunotia* Ehrenberg (2.5 %) and *Stauroneis* Ehrenberg genus (13 %), closely related to the taxa identified by LANGE-BERTALOT et al. (2003) in Sardinia, as *Eunotia sardiniensis* Lange-Bertalot, Cavacini, Tagliaventi & Alfinito and *Stauroneis reichardtii* Lange-Bertalot, Cavacini, Tagliaventi & Alfinito

### Comparison with related taxa

The most similar taxa to *Pinnularia baetica* was *Pinnularia atlasi* and but it was found with more differences *Pinnularia infirma*. (Table 2) The first taxon in this comparison is *P. atlasi* (Figs. 17-31, 53-58). Data and images of *P. atlasi* collected for this work were collected from the locations (Locotype) where DARLEY (1990) described this species only based on LM observations. This study revealed the presence of spines in the populations of Morocco as well as in the population of Sardinia, according to the ultrastructure study performed by LANGE-BERTALOT et al. (2003).

**Table 2.**
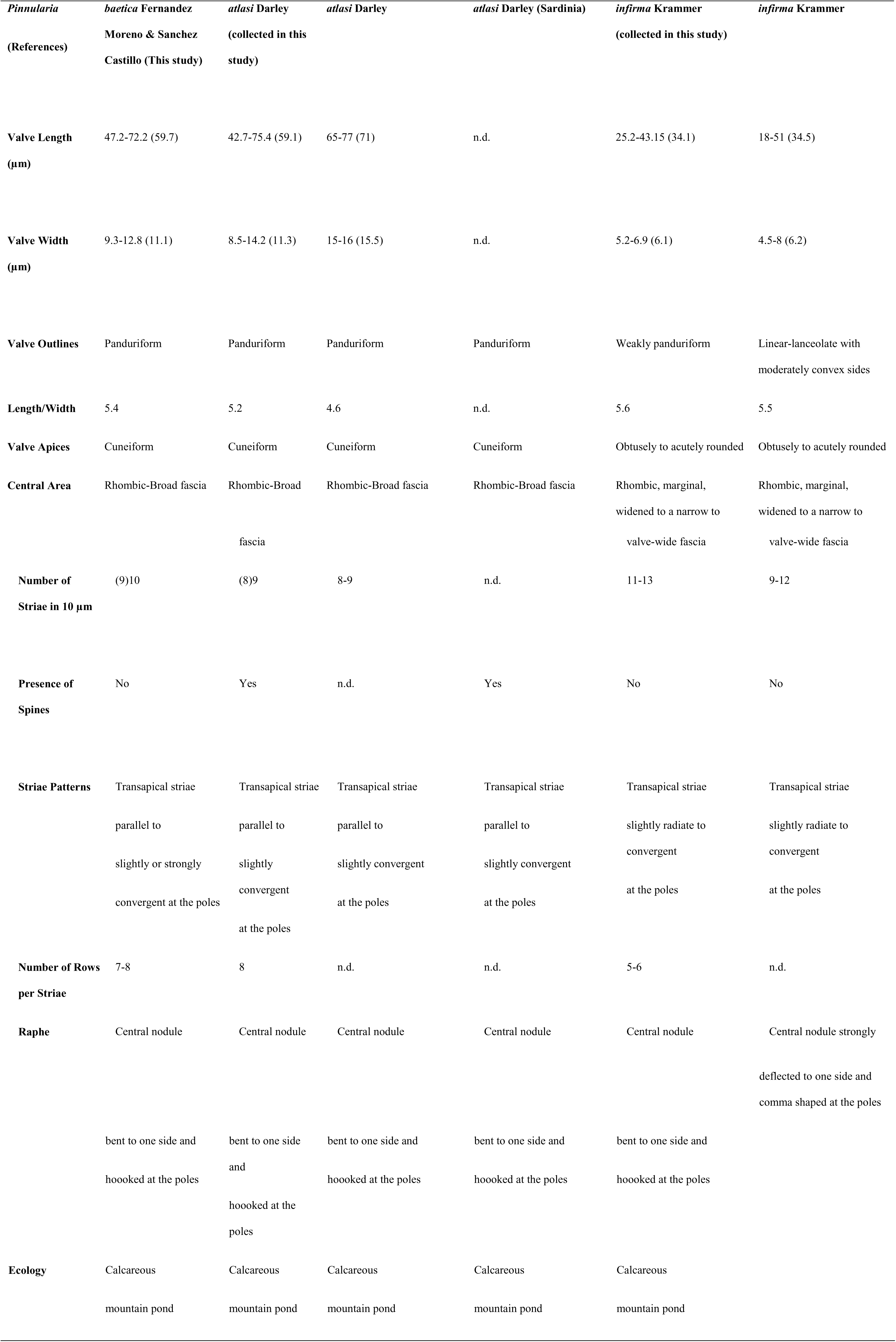
Comparison between *Pinnularia baetica* and the morphologically and ecologically most similar taxa (data from Sardinia comes from the literature)

**Fig. 17-31.**
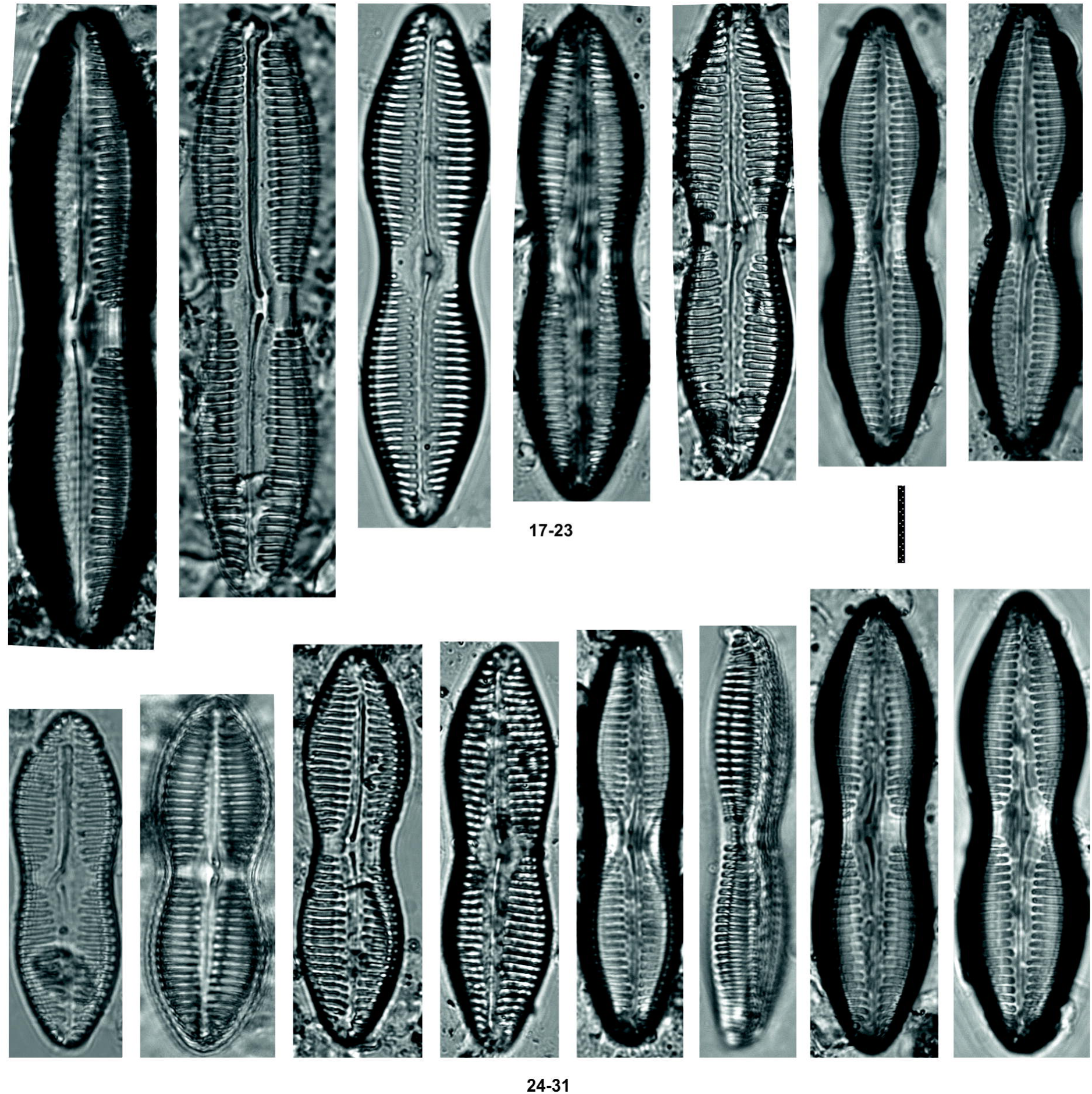
*Pinnularia atlasi* DARLEY. Light micrographs of the type population collected in Morocco. Scale bar 10 μm.

**Fig. 32-46.**
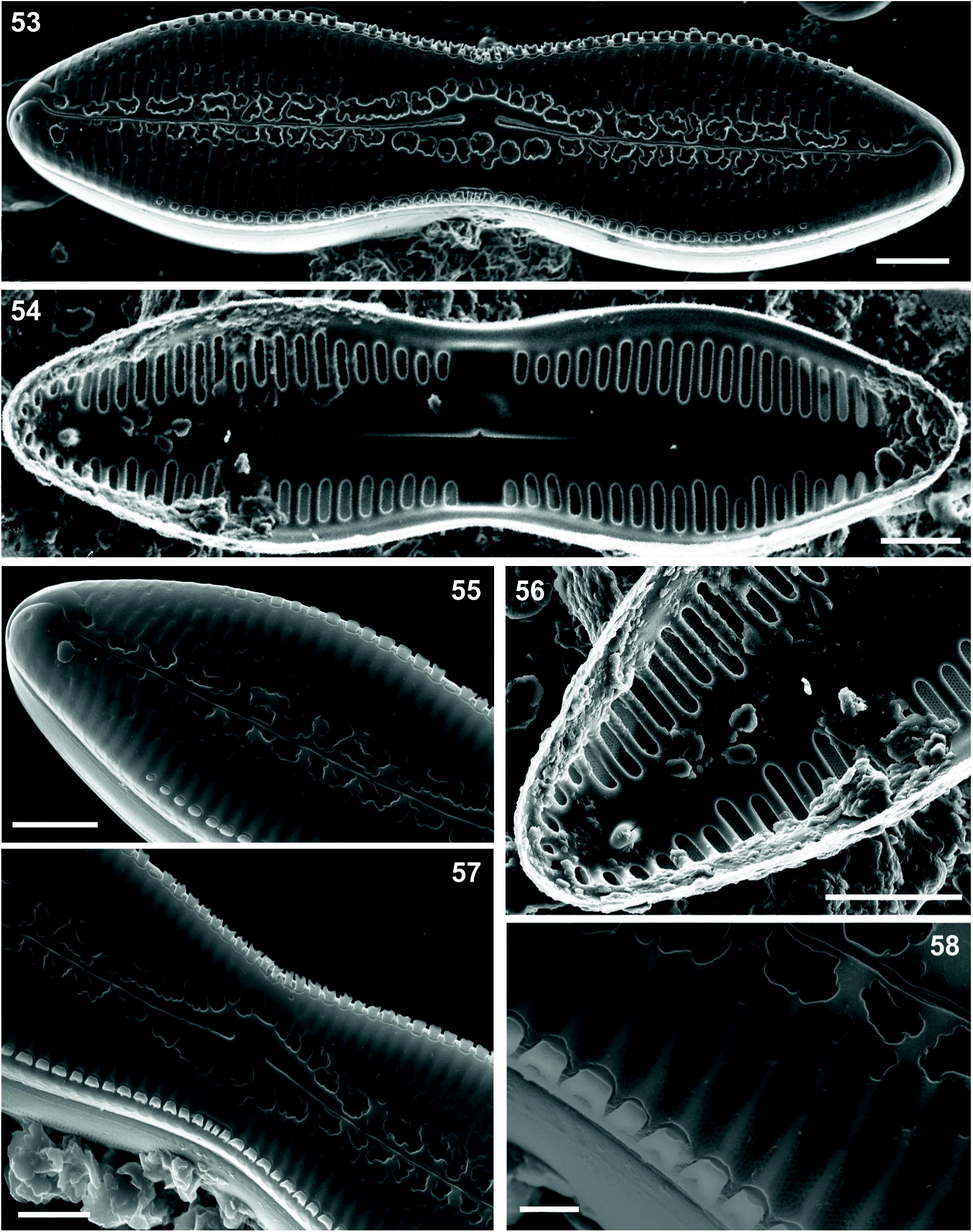
*Pinnularia infirma* KRAMMER. Light micrographs of the type population collected in Ouiouane, Morocco in this study. Scale bar 10 μm

**Fig. 47-52.**
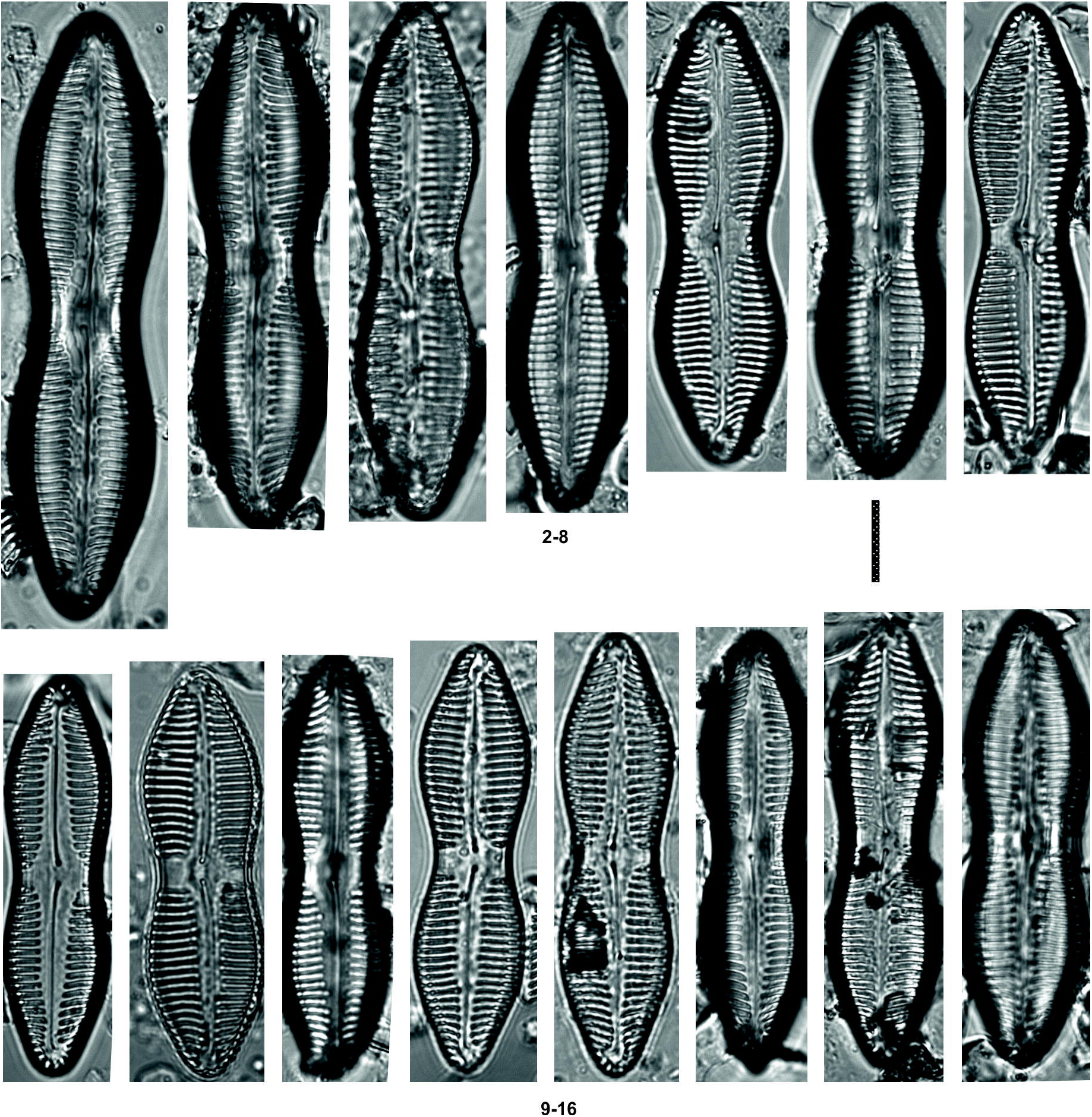
*Pinnularia baetica* sp.nov. Scanning electron micrographs of the type population sampled in Laguna Seca. (47) external valvar view; (48) internal valvar view; (49) external view of the apex, showing the distal raphe hooked endings; (50) internal view of the apex, showing the distal raphe endings terminating on small helictoglossae; (51) external central nodule bent on one side; (52) rows of each alveoli. Scale bars 5 μm (47-49,51); 1 μm (50,52).

This character is rare in the genus (ROUND et al. 1990) and different models have been observed. Among the species with the greatest similarity to *P. atlasi*, in terms of this character, there is *Pinnularia spinea* LANGE-BERTALOT et al. (2003). with the same type of spines and similar sequence of size structure (although it has several rows of spines, being the external smaller) different from other spine-carrying species of the genus *Pinnularia, Pinnularia sofiae* VAN DEN VIJVER et LECOHUMODEL (VAN DE VIJVER et al. 2008), *Pinnularia catenaborealis* (PINSEEL et al. 2016)., *Pinnularia gemella* Van de Vijver in Van de Vijver et al.

In the collected material from the two locations of Morocco, there are little differences in valve length (42.7-75.4 µm), width (8.5-14.2 µm) but the same number of striae 8/9, in 10 µm when compared with the original description of *P. atlasi* from Darley (1990): 65-77 in length, 15-16 in width, 8-9 striae in 10 µm.

The main differences between P.atlasi collected in this study and P.baetica is the valve margin covered over almost its entire length by well-developed spines located in the valve margin. The central spines are relatively large (approx. length/width: 1 µm x 1 µm) with bifurcated margins (Fig 58), while those located towards the ends of the valve are smaller and have blunt margins (Figs 55). They are clearly aligned in a single marginal row between the valve and mantle, progressively decreasing in diameter from the centre to the apex of the valve, even disappearing (Figs 53, 55).

**Fig. 53-58.**
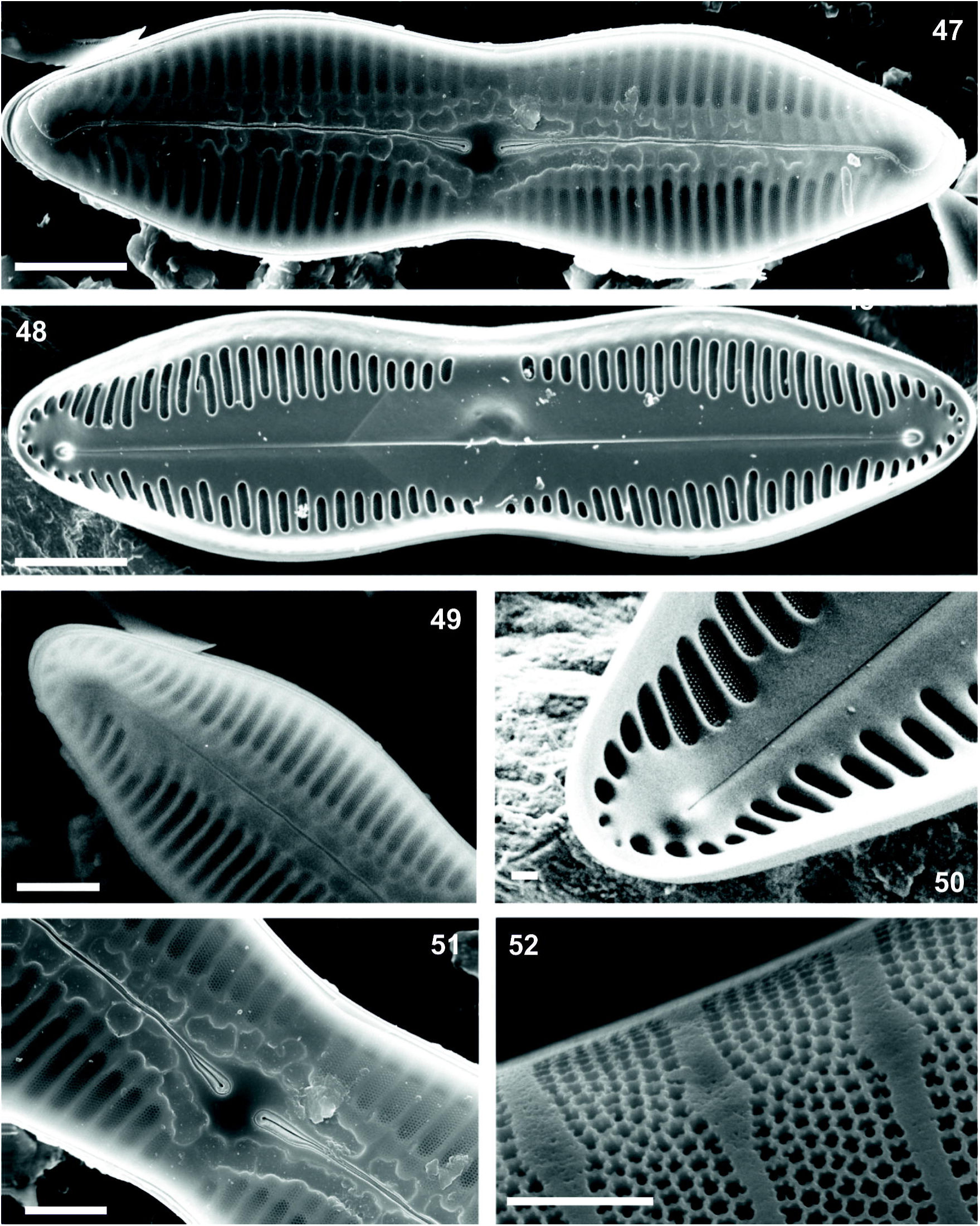
*Pinnularia atlasi* DARLEY. Scanning electron micrographs of the type population collected in Morocco. (53) external valvar view; (54) internal valvar view;(55) external view of the apex, showing the distal raphe hooked endings; (56) internal view of the apex, showing the distal raphe endings terminating on small helictoglossae; (57) external central nodule bent on one side and spines in marginal area; (58) external rows of each alveoli. Scale bars 5 μm (53-57); 1μm (58).

**Fig. 59-64.**
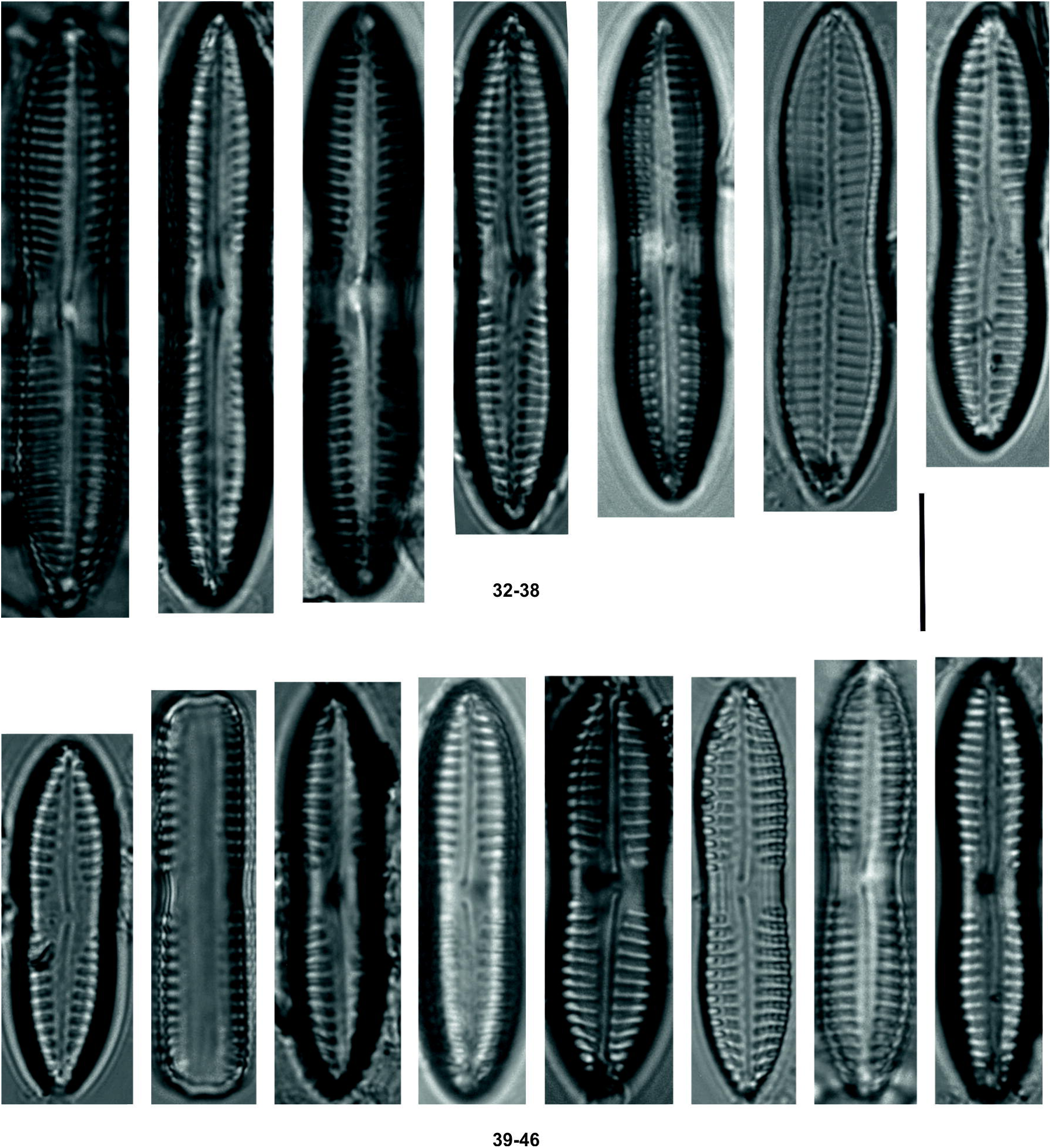
*Pinnularia infirma* KRAMMER. Scanning electron micrographs of the population collected in Ouiouane, Morocco. (59) external valvar view; (60) internal valvar view; (61) external view of the apex, showing the distal raphe hooked endings; (62) external central nodule bent on one side; (63) external rows of each alveoli; (64) Girdle view. Scale bars 5 μm (59-61); 1 μm (62-63); 10 μm (64).

The presence of spines in *P. atlasi* is possible to observe in some views of the frustule in LM (Fig 65) like it is said in LANGE-BERTALOT et al. (2003) (LM: Plate 88: Fig 3).

**Fig. 65.**
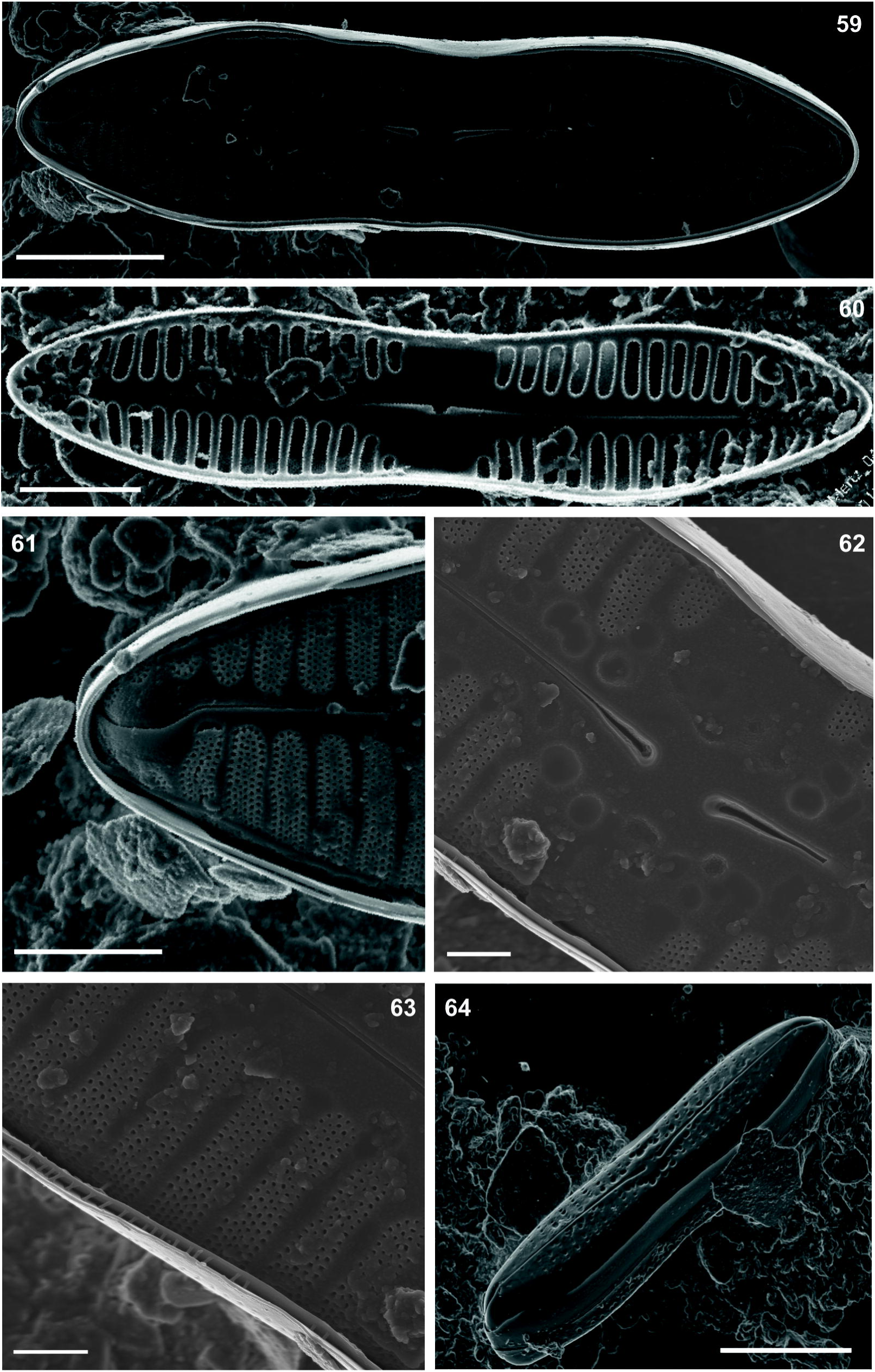
*Pinnularia atlasi* DARLEY from Morocco. Light micrograph in girdle view.Showing the spines (see arrow) defined in the middle of the concave part of the lower valve.

*P. baetica* shows more convergent striae in the apices (Figs 47, 48), but the most evident characteristic is the absence of spines on the valvar margin (Figs 47, 51). Small differences in the number of striae, a bit less in *P. atlasi* (8-9) than in *P. baetica* (9-10). This is because of the fact that in *P. atlasi* the alveolus had 8 rows of areolae, meanwhile in *P. baetica* just 7-8 rows of areolae were found. The length/width ratio of *P. baetica* (5.4) is similar when compared with the same in *P. atlasi* (5.2).

*P. infirma* KRAMMER & LANGE-BERTALOT (1985) a third taxon used for this comparison was for the first time described in central Europe, in a locality at 250 m of altitude (Franken, Germany). *P. infirma* which was also identified in Morocco, in our study, is referenced in many bibliographic references from different places, such as Germany (HUSTEDT 1930; LUDWIG & SCHNITTLER 1996; KRAMMER & LANGE BERTALOT 1985), Great Britain (WHITTON et al. 2003), Macedonia (LEVKOV et al. 2005), Russia (STENINA & PATOVA 2007) and Sardinia (KRAMMER 2000; LANGE-BERTALOT et al. 2003). No ultrastructural data are provided and no reference to the presence of spines is given.

In this study, this taxon is characterized by its narrow central area, length x width 25.2-43.15 x 5.2-6.9 µm and 11-13 striae in 10 μm. In the same sample from Ouiouane ephemeral pond (not found in Guedrouz), the analysis of *P. infirma* could be confused with *P. atlasi*. Neverthless, *P. infirma* differs sufficiently from *P. atlasi*, as well from *P. baetica* on the valve outline.

*P. infirma* had a slight panduriform valve instead of the more panduriform of *P. baetica* furthermore, the length/width relationship is higher in *P. infirma* (5.6) than in *P. baetica* (5.4). *P. infirma* has obtusely to acutely poles, smaller size and the number of rows in each alveolus is lower (5-6) than in *P. baetica* (7-8).

The interesting distribution of the species on this study (north of Morocco, South of Spain and Sardinia) is known for the succession of continuous geological events from the era of the Tertiary that caused the connection and fragmentation of territories between Sardinia and northern Africa through the Balearic Islands and southern Europe (AZZAROLI 1981).

These species, have biogeographical (South Morocco, Spain, Islands like Sardinia) similarities with other endemical organisms from the circumediterranean region such plants (NIETO-FELINER 2014; ROSELLÓ 2013), and other aquatic groups with a high rate of endemicity in the Mediterranean (TIERNO DE FIGUEROA et al. 2013). A better knowledge of the diatom communities in the south of europe could add more information about the biological history in this Mediterranean region.

## Acknowledgements

We are indebted to the Excellence Research Project (RNM-7033) for financial support for the field work in Morocco. To Dr. Z. Letkov who kindly provided us with *P. infirma* material from their collections in Macedonia. To the General Directorate for the Environment of the Andalusian Government who supported the first phase of the “Phycological Flora of Andalusia”. To Hydraena S.L.L. and DNOTA Medio Ambiente for facilitating sampling. To Cristina Delgado and Salomé Almeida for the revision of this article. We are very thankful to Antonio, Miguel, Tite, Javi, Jesus, Sonia and Greta for helping us on the field work for the sampling in Morocco.

